# Selective Disruption of Mutant TP63 Alleles Restores Corneal Epithelial proliferation in EEC Syndrome

**DOI:** 10.64898/2026.02.09.704349

**Authors:** Giulia Masi, Gualtiero Alvisi, Patrizia Nespeca, Anna Demarinis, Chiara Frasson, Luisa Barzon, Vanessa Barbaro, Stefano Ferrari, Giorgio Palu’, Enzo Di Iorio, Marta Trevisan

## Abstract

Ectrodactyly–Ectodermal Dysplasia–Cleft Lip/Palate (EEC) syndrome is a rare disorder caused by dominant-negative mutations in the TP63 gene, frequently leading to limbal stem cell deficiency (LSCD) and progressive corneal degeneration. Current therapeutic strategies are limited, primarily due to impaired epithelial renewal and poor proliferative capacity of patient-derived cells. We have recently shown that decreasing the expression of the mutated allele by means of siRNA-mediated silencing can restore epithelial cell proliferation. However, the clinical utility of this approach is hindered by the presence of different TP63 mutations causing EEC syndrome, and the need for continuous siRNA administration to achieve sustained gene silencing. To address these challenges, we employed a CRISPR/Cas9-based genome editing strategy to disrupt mutant TP63 alleles in human induced pluripotent stem cells (hiPSCs) derived from EEC patients carrying R279H and R304Q mutations. Targeted editing of exon 6 induced frameshift mutations that activated nonsense-mediated mRNA decay, leading to a significant reduction in mutant transcript levels. Edited hiPSC-derived corneal epithelial cells exhibited improved cell proliferation compared to unedited isogenic controls. These findings demonstrate the feasibility and therapeutic potential of allele-specific genome editing to correct TP63-associated epithelial defects in EEC syndrome paving the way toward future regenerative therapies for TP63-related corneal diseases.

## Introduction

Ectrodactyly-Ectodermal Dysplasia-Cleft Lip/Palate (EEC) syndrome is a rare autosomal dominant disorder caused by heterozygous mutations in the TP63 gene, which plays a central role in epithelial development and maintenance^1^. The clinical triad includes ectrodactyly (skeletal abnormalities of hands and feet), ectodermal dysplasia (affecting skin, appendages, and teeth), and orofacial clefts such as cleft lip and/or palate ^2,3^. Ocular involvement is frequent, primarily affecting the corneal epithelium whose maintenance relies on limbal stem cells (LSCs), located in the limbus. The limbus supports epithelial renewal and acts as a physical barrier against conjunctival invasion. Loss or dysfunction of LSCs in EEC patients leads to impaired corneal regeneration, conjunctivalisation and progressive vision loss^1^.

At the molecular level, EEC syndrome is caused by missense mutations in the DNA-binding domain of TP63, located on chromosome 3q28. TP63 produces multiple isoforms through alternative promoters and splicing^4^. ΔNp63α is the primary isoform expressed in basal epithelial cells and is essential for progenitor maintenance^5^. These mutations impair p63 transcriptional activity and exert dominant-negative effects, thus disrupting normal epithelial function^6^. Currently, no definitive treatment exists for the ocular complications of EEC syndrome. We previously demonstrated that siRNA-mediated allele-specific silencing of mutant TP63 mRNA can rescue epithelial defects and restore normal epithelial stratification in oral mucosa epithelial stem cells (OMESCs) derived from EEC patients^7^. Despite this approach is promising, its clinical utility is hampered by the need of a specific siRNA for each of the more than 10 different TP63 mutations, and continuous administration. Importantly both such limitations would be resolved by genome editing of the mutated allele targeting an upstream exon. *Ex vivo* cell therapies, based on gene editing of patient-derived cells are currently under investigation. However, the absence of residual limbal stem cells in many EEC patients beyond the age of 20–25 significantly limits the feasibility of transplanting genetically corrected autologous limbal grafts^1^, and the utility of patients-derived OMESCs is significantly impaired by premature senescence and diminished regenerative potential due to TP63 mutations^7^. In this study, we investigated induced pluripotent stem cells (hiPSCs) derived from EEC patients as an alternative cell source for gene editing purposes. Our results demonstrate that CRISPR–Cas9 knockout of the mutant TP63 allele in EEC patient–derived hiPSCs re-establishes wild-type expression and restores the proliferative potential of corneal epithelial–like cells. Overall, our findings provide the first evidence that allele-specific genome editing can restore the proliferation of corneal epithelial cells in EEC syndrome, paving the way for translational development.

## Material and Methods

### Cell cultures

hiPSCs were previously generated form OMESCs of an healthy donor and two EEC patients harboring the mutations R279H and R304Q, as previously described ^8,9,10,11^. hiPSCs were maintained in culture in Geltrex (Gibco, Thermo Fisher Scientific, Waltham, Massachusetts, USA)-coated plates in mTeSR-1 medium (Stemcell Technologies, Vancouver, BC, Canada) and passaged with Accutase (Invitrogen, Thermo Fisher Scientific). Human embryonic kidney 293A cells (HEK-293A, Thermo Fisher Scientic) were grown in Dulbecco’s Modified Eagle’s Medium (DMEM, GIBCO) supplemented with 10% Fetal Bovine Serum (FBS), 1% GlutaMAX™, 1% Penicillin/Streptomycin (all from GIBCO).

### Plasmids

U6gRNA-Cas9-2A-GFP plasmids were obtained from Sigma Aldrich (St. Louis, Missouri, USA); guide RNA (gRNA) sequences are the following (PAM sequence in square brackets):

gRNA-1:[CCC]TCCTAGTCATTTGATTCGA;

gRNA-2:[CCT]CCTAGTCATTTGATTCGAG;

gRNA-3: [CCT]AGTCATTTGATTCGAGTAG.

### Transfection experiments

Transfection of HEK-293A (5×10^5^) was carried out by mixing 500 ng of CRISPR/Cas9-2A-GFP plasmids with 2.5 μl of Lipofectamine®2000 (Invitrogen, Thermo Fisher Scientific). Forty-eight hours post transfection, cells were harvested for further analysis. One million hiPSCs (seeded in which format) were transfected with the CRISPR/Cas9 plasmids combination (1.5 μg of the three sgRNA plasmids and 1.5 μg of pCas9-GFP plasmid) by using 3.75 μl Lipofectamine®3000 (Invitrogen, Thermo Fisher Scientific).

### Surveyor assay

Genomic DNA (gDNA) isolated from transfected HEK-293A cells and hiPSCs (QIAampDNA kit, Qiagen, Hilden, Germany) was used to amplify the region in exon 6 of TP63 gene (A list of primers used is available in Table S1). Heteroduplex formation and digestion were performed following Surveyor Mutation Detection kit’s manufacturer recommendation (Integrated DNA technologies, Coralville, Iowa, USA). A not-digested control was also included. The final products were then electrophoretically separated on a 10% polyacrylamide gel in 1X TBE. Subsequently, the gel was stained with GelRed (Sigma Aldrich) in 1X TBE for 15 minutes and washed twice in TBE. Cleavage efficiency was determined using ImageJ software^12^ by measuring the optical density (volume of the area under the curve) of uncleaved and cleaved PCR products.

The fraction of cleaved PCR product was calculated using the formula:

*(Σ area of cleaved bands/ Σ area of cleaved and uncleaved bands)*.

The cutting efficiency was estimated using the equation:

*% cutting efficiency = 100 x [1 -(1-fraction cleaved)*^*1/2*^].

### Cell sorting of transfected hiPSCs

Seventy-two hours post-transfection, hiPSCs were sorted with a FACSAria™ III Cell Sorter (BD Bioscience) to select green fluorescent protein (GFP) expressing (GFP+) cells. hiPSCs were detached with Accutase and passed through a Falcon® 40μm Cell Strainer (Corning, NY, USA). Cells were re-suspended in a sorting solution (PBS supplemented with 1% FBS, 1% penicillin/streptomycin) containing 10 μM Y-27632 (Sigma Aldrich) and GFP+ cells were collected in a tube containing mTeSR1 supplemented with 10 μM Y-27632. Sorted cells were plated on Matrigel-coated 10-cm^2^ dishes at densities ranging from 3×10^4^ to 5×10^4^ cells per plate. The medium was then changed every other day while cells recovered, and then daily once multicellular colonies had formed.

### hiPSC clones expansion and screening

Approximately 10 days after plating, individual, single cell–derived colonies were selected and picked. Small clumps were generated and gently transferred into a Geltrex-coated 96-well plate with mTesR-1 and 10 μM Y-27632. Clones were then passaged with Accutase in two different Matrigel-coated 96-well plates: one plate was used for gDNA isolation, while the other one for cell expansion^13^. gDNA was isolated from cells on the 96-well plate using an automated DNA extractor (MagNA Pure 96 System, Roche, Switzerland). PCR amplification was performed on isolated gDNA using primers reported in Table S1. Genome edited clones were confirmed by direct Sanger sequencing of PCR products. The final editing outcome was predicted using the MutationTaster online software (*www*.*mutationtaster*.*org*^14^).

### RNA extraction of differentiated hiPSCs

Differentiation of EEC-derived hiPSCs into corneal epithelial-like cells was performed as previously described^8^. Cells on day 10 of differentiation were collected by trypsinization and RNA was extracted using RNeasy kit (Qiagen) following manufacturer’s instructions.

### TA cloning analysis of edited clones

TP63 cDNA fragments were amplified with an AmpliTaq Gold DNA Polymerase (Applied Biosystems, Thermo Fisher Scientific) to promote A-tailing. The primers used are reported in Table S1. Amplicons were ligated into pGEM-T Easy Vector (Promega, Madison, Wisconsin, USA) following manufacturer’s protocol and transformed in DH5α cells. Transformed bacteria were spread on selective plates of Luria-Bertani (LB) agar containing 100 μg/mL ampicillin (both from Sigma Aldrich). The day after, single white colonies were picked and directly pipet into the PCR mixture. Amplicons were purified through ExoSAP-IT™ PCR Product Cleanup Reagent (Applied Biosystems). Sanger sequencing reaction was performed using the BigDye® Terminator v3.1 Cycle Sequencing Kit (Applied Biosystems). Following sequencing reaction, amplicons were purified through precipitation with 3M sodium acetate and 96% ethanol. Subsequently, they were loaded on an ABI PRISM 3130xl Genetic Analyzer (Thermo Fisher Scientific).

### Off-target analysis

The CRISPR design tool (Sigma) was employed to predict potential off-target sites scattered throughout the entire genome. The identified potential undesired editing sites were amplified and directly sequenced, as described above, with specific primer pairs (Table S1). Sequences in FASTA format were aligned to human genome by employing the multiple sequence alignment ClustalW software (*2*.*1 version, https://www.genome.jp/tools-bin/clustalw)^15^.*

### Real Time PCR analysis

cDNAs from EEC-derived hiPSCs and differentiating clones were amplified in an ABI PRISM® 7000 Sequence Detection System (Applied Biosystems). Glyceraldehyde-3-phosphate dehydrogenase (GAPDH) expression was used to normalize the results between samples. Primers and TaqMan probes sequences used are reported in Table S1.

### MTT assay

Edited and isogenic hiPSCs were differentiated into corneal epithelial-like cells in 24-well plates following a 14-day differentiation protocol^8^. The wt-hiPSC line was included as control. At the end of the differentiation period, cell viability was assessed using the 3-(4,5-dimethylthiazol-2-yl)-2,5-diphenyl tetrazolium bromide assay (MTT, Sigma-Aldrich) according to the manufacturer’s instructions. After incubation with MTT reagent, formazan crystals were solubilized, and absorbance was measured at 570 nm using a microplate reader (Varioskan Lux, Thermo scientific).

### Statistical analysis

Data analysis was performed with GraphPad Prism version 10.4.1 (GraphPad, San Diego California USA). All graphs and applied statistical tests are detailed in figure legends.

## Results

### TP63 Exon 6-targeting gRNAs and CRISPR/Cas9 efficiently induce indels in hiPSCs

hiPSCs from healthy donors and EEC patients harboring the R279H and R304Q TP63 mutations were previously generated and characterized^8–10^. To induce knockout of the mutant allele, we designed three guide RNAs (gRNAs) targeting exon 6 of TP63 (Fig. 1A-B). To assess editing efficiency, we first co-transfected HEK-293A cells with a plasmid encoding a *Streptococcus pyogenes* Cas9 fused to a GFP and the gRNAs, either individually or combined. Editing efficiency was evaluated by Surveyor nuclease assay on pooled transfected cells, revealing comparable efficiencies for all three gRNAs, with an average indel frequency of approximately 29% (Fig. 1C). We next tested the editing system in hiPSCs derived from a wild-type donor^9^, combining the three gRNAs-Cas9 plasmids to enhance on-target efficiency^16^. After transfection, a mixed population of edited and unedited cells was analyzed by Surveyor assay, showing roughly half the editing efficiency observed in HEK-293A cells (Fig. 1D). To improve genome-editing efficiency, wt-hiPSCs, R279H-hiPSCs and R304Q-hiPSCS, were sorted for high GFP expression three days after transfection. Sorted cells were then plated at low density, allowing the formation of individual colonies, which were subsequently isolated and expanded (Fig. 2A). A total of 234 clones were screened for Non-Homologous End Joining (NHEJ) events. Genomic DNA was extracted from each clone, and a ∼400 bp region of TP63 exon 6 encompassing the CRISPR target site (Fig. 1B, light blue arrows) was amplified and sequenced. Analysis of electropherograms revealed insertions and deletions (indels) near the cleavage site. Indels were detected in 15 of 43 wt-hiPSC clones, 41 of 192 R279H clones, and 24 of 95 R304Q clones, yielding an average NHEJ efficiency of 27% overall (34.8% in wt-, 21.3% in R279H-, and 25.2% in R304Q hiPSCs) (Fig. 2B-C). Most indels consisted of deletions, though insertions of single or paired nucleotides were also observed (Fig. 2D). Collectively, these results demonstrate that our editing strategy efficiently targets TP63 exon 6 with comparable efficiencies across hiPSCs derived from different donors.

**Fig. 1.**
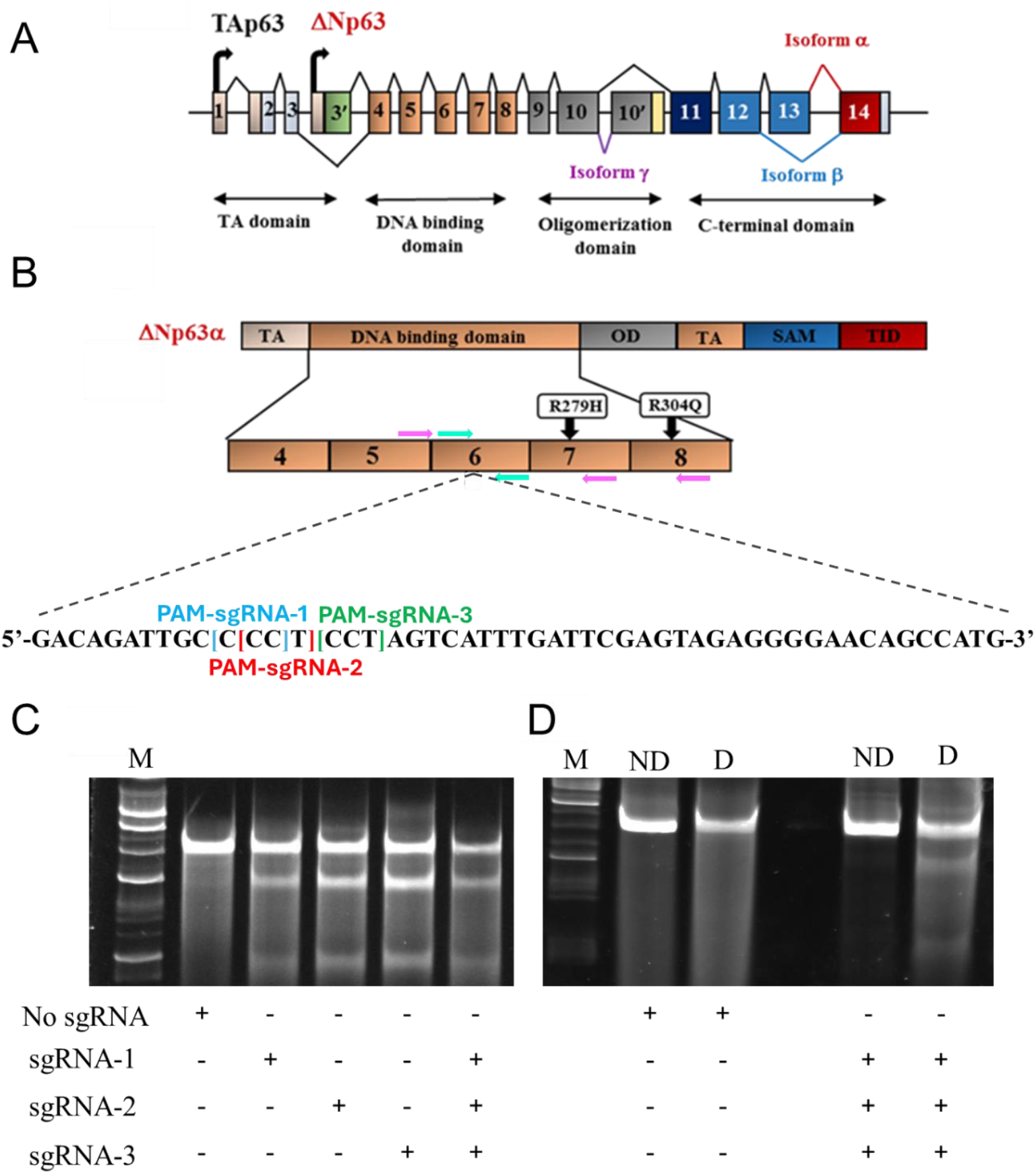
TP63-exon-6-targeting gRNAs and CRISPR/Cas9 display a comparable targeting efficiency. **A** Graphical scheme of TP63 gene. **B** Graphical scheme of the genome editing approach. Colored squares represent different ΔNp63α domains: TA: transactivation domain; OD: oligomerization domain; SAM: sterile alpha motif; TID: transactivation inhibitory domain. Orange squares represent exons of the DNA binding domain. Nucleotide sequence represents Exon 6 target sequence of sgRNAs. Square brackets represent the PAM sequences matched with the three guide RNAs (gRNAs) used. EEC-patient mutations are represented in exon 7 and 8 (R279H and R304Q, respectively). Light blue arrows represent forward and reverse primers used to amplify the CRISPR/Cas9 target region in exon 6 on genomic DNA; purple arrows represent forward and the two reverse primers used to discriminate alleles carrying edits or mutations on cDNAs. **C-D** Surveyor assay on HEK-293A cells (C) and wild-type hiPSCs (D) transfected with the single gRNAs or in combination (D, digested; ND, not digested; C-, negative control; M, molecular marker VIII).

**Fig. 2.**
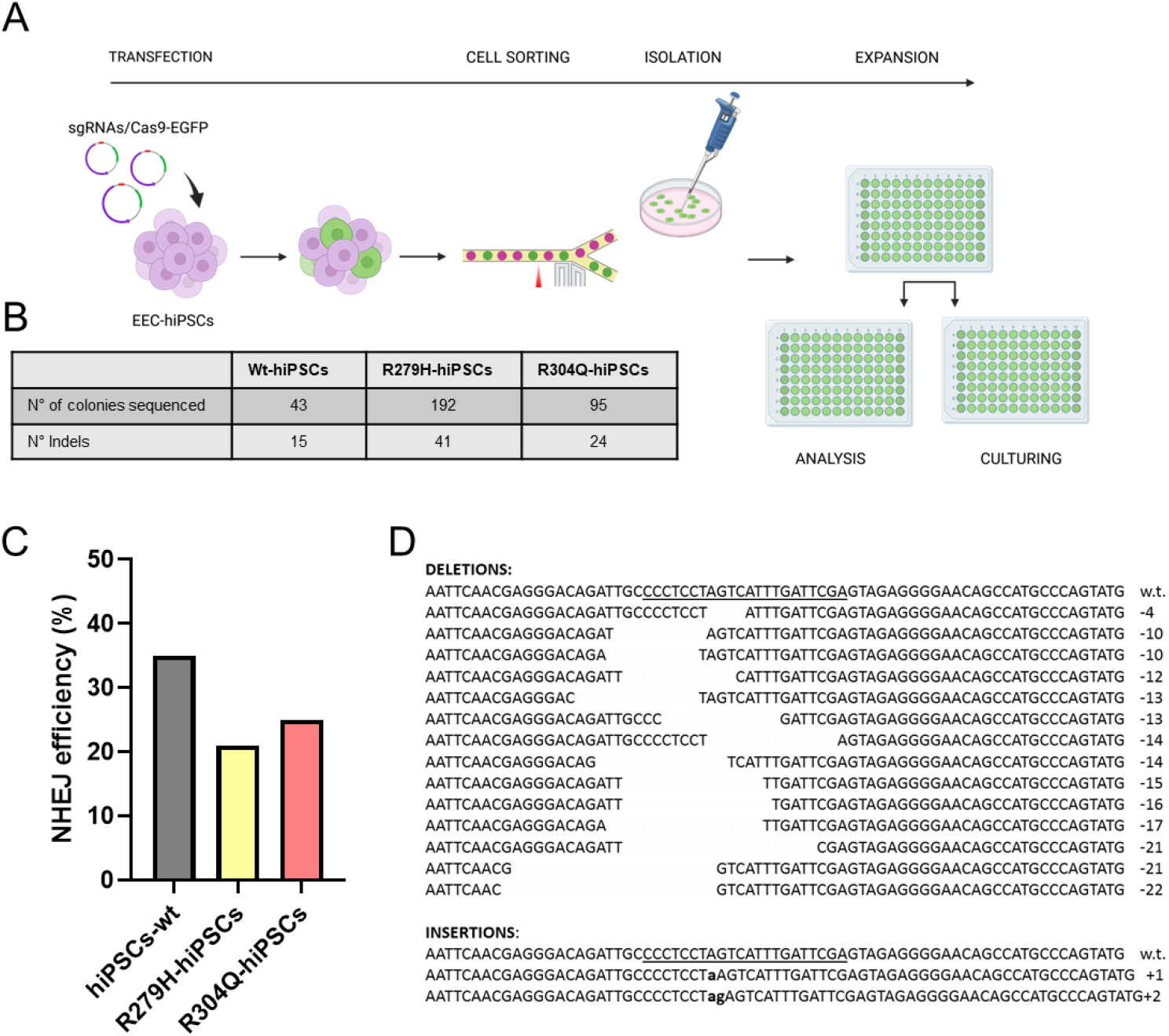
TP63-exon-6-targeting gRNAs and CRISPR/Cas9 display a similar targeting efficiency in cells originating from different donors. **A** Scheme of the genome editing protocol used. hiPSCs were transfected with lipofectamine 3000 and sorted for the GFP signal. Sorted cells were plated at low density and allowed to grow in single colonies. Once grown, single colonies were picked and plated on 96 well plates and passaged for subsequent analysis and subculturing. This schematic was created with BioRender.com. **B** Number of indels reported per hiPSC cell line. **C** Bar graphs showing the total percentage of efficiency of Non-homologous end joining (NHEJ) in edited clones. **D** Representative insertions/deletions (indels)-containing sequences compared to the wild-type target region.

### Edited EEC-hiPSC clones retain pluripotency

For each editing experiment, a total of fourteen clones of R279H- and R304Q-hiPSCs (n = 7 for each line) harboring indels (Fig. S1) were further cultured and analyzed to predict whether the introduced modifications induced a frameshift resulting in a premature stop codon, likely triggering nonsense-mediated mRNA decay (NMD)^17^. To this end, the *MutationTaster* tool (https://www.mutationtaster.org/) was employed for in silico prediction. Due to the large intronic distance between the editing site and the endogenous mutation, direct determination of the allele bearing the editing event at the genomic DNA level was not feasible. Therefore, this allelic discrimination was performed at the mRNA level. However, since pluripotent stem cells do not express TP63 transcripts, hiPSC clones were differentiated into corneal epithelial-like cells^8^, to enable TP63 expression and subsequent analysis. Total RNA was extracted from differentiated cells, and TA cloning was performed on PCR products amplified with primers spanning both the editing and mutation sites (Fig. 1B, purple arrows). For both R279H and R304Q analyzed clones, approximately half showed a mixture of edited and non-edited sequences on both the wt and mutant allele, at different ratios, suggesting that these colonies likely originated from a heterogenous population of edited cells rather than single-cell-derived clones and therefore require further purification (Table S2). In contrast, the remaining clones presented the modifications only in one allele (Table S2). Among these, for R279H-hiPSCs, a 14 bp deletion (NM_003722: c.772_785del14; I258Sfs*29) was identified in the mutant allele of clone R279H-hiPSCs^wt+/R279H−^ (Fig. 3A). Similarly, in R304Q-hiPSCs, an insertion of an adenine (NM_003722: c.784_785insA; S262Kfs*30) was detected in the mutant allele of clone R304Q-hiPSCs^wt+/R304Q−^ (Fig. 3A). Both mutations were predicted to cause a premature stop codon and NMD. Importantly, the selected clones retained expression of pluripotency markers OCT4, SOX2, and NANOG (Fig. 3B-C) and showed no detectable off-target effects as assessed by targeted sequencing (Figs. S2-S12) and were therefore used for further analysis.

**Fig. 3.**
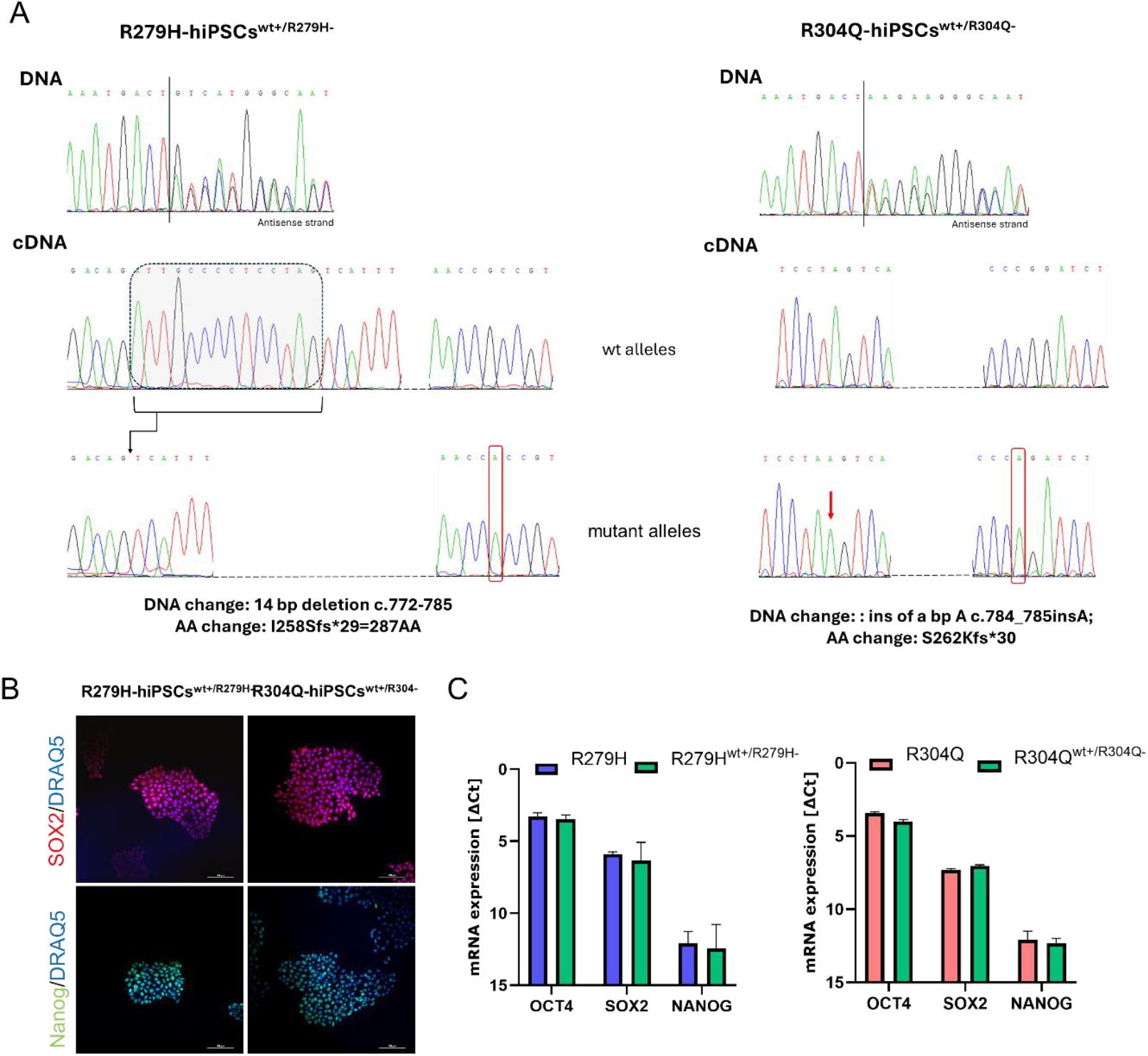
Edited EEC-derived hiPSC clones maintain their pluripotent potential. **A** Electropherograms showing the types of modifications detected in selected edited EEC-derived hiPSC clones. R279H-hiPSCs^wt+/R279H-^ display a frameshift in the genomic (DNA) sequence caused by a 14bp deletion in the mutant allele (cDNA). R304Q-hiPSCs^wt+/R304Q-^ show in the DNA sequence a frameshift exerted by an insertion of an A (red arrow) in the mutant allele (cDNA). Red rectangles highlight the G>A nucleotide change present in the mutant alleles of both R304Q and R279H. **B** Representative images of the expression of SOX2 and Nanog in edited R279H-hiPSCs^wt+/R279H-^ and R304Q-hiPSCs^wt+/R304Q-^. Nuclei were counterstained with DRAQ5. Scale bars: 100 µM. **C** Bar graph showing *OCT4, SOX2* and *Nanog* mRNA levels (expressed as delta Ct calculated over the *GAPDH* housekeeping gene) in selected edited EEC-hiPSCs. Data show the mean ± SEM of n = 3 independent experiments.

### Edited EEC-hiPSC clones show activation of NMD and reduction of mutant allele expression

To confirm the successful knockout of the mutated allele, we evaluated the activation of the NMD pathway in edited clones R279H-hiPSCs^wt+/R279H−^ and R304Q-hiPSCs^wt+/R304Q−^. Both edited hiPSC clones, along with their respective isogenic controls, were differentiated into corneal epithelial-like cells, and total RNA was extracted after 15 days of differentiation. PCR amplification was performed using primers spanning both the editing and mutation sites. The PCR products were then subjected to TA cloning to separate the two alleles, enabling assessment of the relative abundance of mRNA bearing the wild-type and EEC mutation containing alleles. One hundred colonies from each clone were analyzed by Sanger sequencing. As expected, both original patient lines showed a predominant expression of the mutated allele over the wild-type^18^ with the R279H-hiPSCs and the R304A-hiPSC line exhibiting 76% and 57% mutated allele frequency, respectively (Fig. 4A). In the edited clones, a marked reduction in the mutant allele was observed: R279H-hiPSCs^wt+/R279H−^ retained a small fraction (∼12%) of the mutated sequence, which was completely undetected in R304Q-hiPSCs^wt+/R304Q^ (Fig. 4A). Therefore, our results suggest that introduction of indels resulting in premature stop codons efficiently triggered NMD and reduced expression of the mutated alleles.

**Fig. 4.**
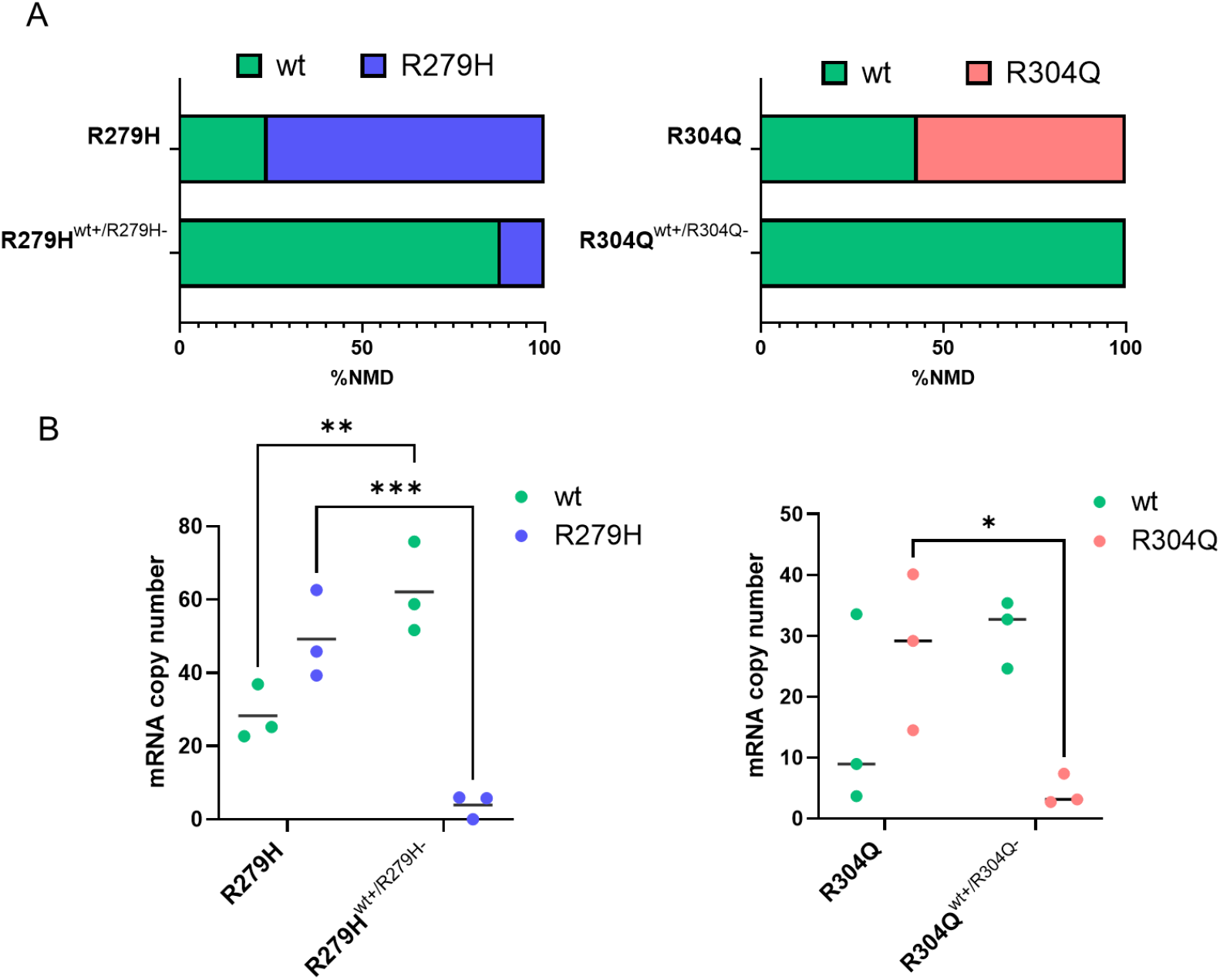
Edited EEC-hiPSC Clones show activation of NMD and reduction of mutant allele expression. **A** Stacked bars showing the activation of the non-sense mediated decay (NMD) pathway expressed as the percentage of sequences containing the wt *versus* mutant allele, obtained from the TA cloning analysis in edited R279H-hiPSCs^wt+/R279H-^ (left panel) and R304Q-hiPSCs^wt+/R304Q-^ (right panel), compared to their isogenic non-edited lines. **B** Dot plots showing the reduction in the expression of mutant alleles in edited R279H-hiPSCs^wt+/R279H-^ (left panel) and R304Q-hiPSCs^wt+/R304Q-^ (right panel), compared to their isogenic non-edited lines, obtained with an allele-specific qPCR. Data are shown as means ± SEM of n ≥ 3 independent experiments, each performed in duplicate, and were analyzed using ordinary two-way ANOVA (*p<0.05, *versus* R304Q-hiPSCs mutated allele; **p<0.01, *versus* R279H-hiPSCs wild-type allele; ***p<0.001, *versus* R279H-hiPSCs wild-type allele).

To corroborate these findings, allele-specific real-time PCR assays, previously optimized for both mutations^19^, were performed. Consistent with the TA cloning results, the original R279H-hiPSCs and R304Q-hiPSCs expressed both wild-type and mutated alleles, with higher expression from the mutant alleles. In contrast, both isogenic edited R279H-hiPSCs^wt+/R279H−^ and R304Q-hiPSCs^wt+/R304Q-−^ lines displayed a significant reduction of mutant allele expression (Fig. 4B). Together, these data demonstrate that the genome editing strategy effectively activates the NMD pathway, resulting in a significant reduction of mutant allele expression in patient-derived hiPSC lines.

### Knock-out of mutant alleles in EEC-derived hiPSCs increases proliferative ability

One of the major challenges in working with patient-derived EEC cells is their severely limited proliferative capacity, primarily due to mutations in TP63, which prevent them from expanding beyond a few passages in culture^5^ and is believed to be reason for ocular manifestations in EEC patients. We therefore evaluated the effect of suppression of mutated allele expression in edited and parental isogenic hiPSC lines. hiPSCs were differentiated into corneal epithelial-like cells, and their proliferation was assessed using an MTT assay. Both original EEC-derived hiPSC lines, R279H-hiPSCs and R304Q-hiPSCs, exhibited significantly reduced cell number compared to the wild-type control (Fig. 5A-B), confirming that epithelial-like cells differentiated from EEC hiPSCs faithfully recapitulate the functional defects observed in primary patient-derived cells. Notably, the genome-edited clones, R279H-hiPSCs^wt+/R279H−^ and R304Q-hiPSCs^wt+/R304Q−^, showed a significant improvement in cell viability compared to their unedited isogenic counterparts (Fig. 5A-B), demonstrating that targeted knockout of the mutant allele effectively restores viability of these cells. Overall, these findings validate the feasibility and efficacy of a CRISPR/Cas9-mediated mutant allele knockout strategy for rescuing cell viability defects in EEC-derived hiPSCs.

**Fig. 5.**
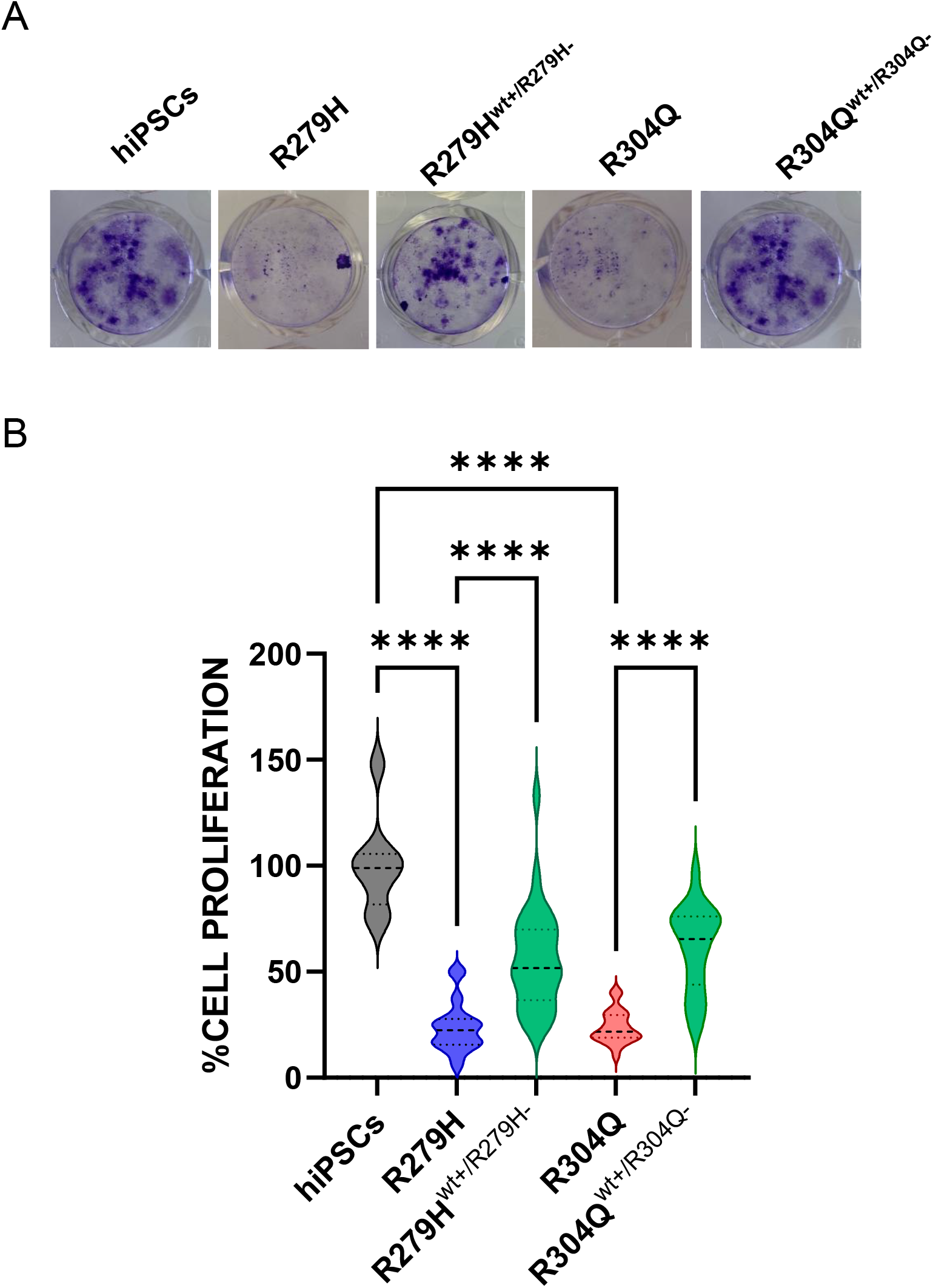
Knock-out of mutant allele in EEC-derived hiPSCs restores cell proliferation. **A** Representative image of cell proliferation assay (MTT) performed on corneal keratinocytes-like cells differentiated from corrected EEC-derived hiPSCs, compared to wt-hiPSCs. **B** Violin plots showing the quantitative comparison of the percentage of cell proliferation in corneal epithelial-like cells differentiated from corrected EEC-derived hiPSCs, compared to isogenic non-corrected lines and wt-hiPSCs. Data represent the percentage of cell proliferation of n ≥ 4 independent experiments, each performed in duplicate. The dashed line represents the median, while the dotted lines represent the quartiles. Data were analyzed with a Kruskal-Wallis test followed by Dunnett’s multiple comparison test. (****p<0.0001, R304Q-hiPSCs and R279H-hiPSCs *versus* wild-type-hiPSCs or edited R279H-hiPSCs^wt+/R279H-^ and R304Q-hiPSCs^wt+/R304Q-^ *versus* R304Q-hiPSCs and R279H-hiPSCs).

## Discussion

In this study, we demonstrated the feasibility and efficacy of a CRISPR/Cas9-based genome editing strategy to knock out mutant TP63 alleles in hiPSCs derived from EEC syndrome patients. Introduction of indels within exon 6 of TP63, which lies upstream of most known pathogenic mutations, resulted in allele-specific reduction of the mutant transcripts, consistent with the activation of NMD^20^. Edited EEC-derived hiPSC clones exhibited markedly improved proliferation ability compared to their isogenic parental lines, upon differentiation into corneal epithelial-like cells. The observed phenotype is unlikely due to off-target effects, since off-target analysis revealed no detectable unintended mutations at predicted *loci*. Therefore, despite unbiased genome-wide approaches are needed to fully rule out rare or structural alterations^21^, our results support the potential therapeutic impact of reducing the expression of the mutant allele to address the ocular symptoms of EEC patients, either at the mRNA^7^ or at the genomic level, as previously shown for other genetic diseases caused by expression of missense mutation alleles, such as Transthyretin Amyloidosis^22^. Importantly, this approach bypasses the key limitations of allele-specific siRNA-mediated silencing of mutant TP63, such as incomplete knockdown, transient effects, and potential off-target repression, previously used to rescue the proliferative defects of EEC patient-derived OMESCs. The specific targeting of exon 6—which lies upstream of the vast majority of EEC-associated mutations—induces a permanent frameshift that ablates production of mutant ΔNp63α, making this strategy broadly applicable to virtually all EEC patient-derived cells. Although low levels of mutant transcripts were still detectable in corneal epithelial–like cells differentiated from the edited hiPSC clones, this residual expression is unlikely to interfere with wild-type p63 function. The truncated proteins generated after exon-6 disruption lack both the DNA-binding domain and the oligomerization domain and are therefore expected to be transcriptionally inert and incapable of exerting dominant-negative effects ^23^. Despite these encouraging results, a few limitations deserve consideration. First, although the genome editing strategy deployed here was efficient and reproducible across different TP63 mutations (R279H and R304Q), the long-term safety and fidelity of CRISPR-based editing must be rigorously assessed, particularly with respect to genomic instability^25,26^. Another limitation relates to the translational applicability of this approach to patient-derived primary cells. Indeed, while genome editing of hiPSCs derived from EEC patients is remarkably easier than editing primary limbal cells or OMESCs, which exhibit severe proliferative impairment due to TP63 dysfunction, their utility for treating ocular diseases is still under investigation. In this context, a recent report described the transplantation of hiPSCs-derived corneal epithelial-like cells as an alternative source for the treatment of limbal stem cell deficiency, suggesting that edited hiPSCs might be similarly used for *ex vivo* transplant of EEC patients^24^. Alternatively, treatment with γ-secretase inhibitors like DAPT has recently been shown to enhance the proliferative capacity of OMESCs derived from EEC patients ^14^, thereby potentially improving the feasibility of genome-editing approaches.

In conclusion our study provides a proof-of-concept for the use of allele-specific genome editing to restore corneal epithelial proliferation in EEC-derived cells. While technical hurdles remain, the strategy outlined here lays the groundwork for future therapeutic applications aimed at correcting TP63-related corneal epithelial pathologies.

## AUTHOR CONTRIBUTIONS

Conceptualization: GM, GA, EDI, MT; Methodology: GM, GA, EDI, MT; Investigation: GM, PN, ADM, MT; Formal analysis: CF (cell sorting); Writing – Original Draft: GM, GA, VB, EDI, MT; Writing – Review & Editing: GA, LB, GP, SF; Supervision: GP, MT, EDI

