## Supplemental Table S1-2; Supplemental Figures S1-12 for "Selective Disruption of Mutant TP63 Alleles Restores Corneal Epithelial proliferation in EEC Syndrome"

**Table S1-S2**

**Supplementary Figures S1-12**

**Supplementary Table 1:** List of oligonucleotides for qPCR used in this study.

| Target | Primer Forward (5'-3') | Primer Reverse (5'-3') |
|---|---|---|
| | **Surveyor analysis** | |
| **P63-exon 6 (gDNA)** | GCCACCAACATCCTGTTCAT | GGGAAAGCATGTGGAGAACA |
| | **TA-cloning analysis** | |
| ***Exon 6-R279H (mRNA)*** | CCTCCTCAGGGAGCTGTTATC | GGACTTGCCCATCTCTGGT |
| ***Exon 6-R304Q (mRNA)*** | CCTCCTCAGGGAGCTGTTATC | CTTCGTACCATCACCGTTCTTT |
| | **Pluripotency analysis** | |
| ***SOX2*** | CCACCTACAGCATGTCCTACTCG | GGGAGGAAGAGGTAACCACAGG |
| ***NANOG*** | GGTCTCGATCTCCTGACCTTGT | GCCTGTAAATCCCAGCTGTTAGG |
| ***OCT4*** | GGTGCCTGCCCTTCTAGGAATGG GGGA | CAAAAACCCTGGCACAAACTC |
| ***GAPDH*** | AGCCACATCGCTCAGACAC | GCCCAATACGACCAAATCC |
| | **Off-target analysis** | |
| **Chr1** | TTCAACCAGGTCACCATCAC | GGTGGTAAAGAGAACACCATCT |
| **Chr2F** | GGGACTGACCAACAGTCTCAA | CTCCCAAAGTGCTGGGATTA |
| **Chr3F** | ATGCCAGGCCTAGAAGTTGA | CAATGCTGAGAACCACTGGA |
| **Chr6F** | GCCCTGCAGAAGGATGTTAT | CTGCATGTGGGTCAAGATCA |
| **Chr8F** | CGGGATTCTACAGAGAAACAGAA | CAGGGCTGCTGAAGTTGATT |
| **Chr10F** | CGGGGCAACCTTACAAAATA | TCTCTTGCATGAGGCTTGAA |
| **Chr12F** | GGCAAGGAAAACTGTCCTGA | TGACAGTGCATCCTCACCTG |
| **Chr14F** | ACCCATTTGGGCTTCTTTTC | GACCTCTCAGGAAAGAAAATTCA |
| **Chr17F** | TTACCCACCATCCTACTACCTC | CGTCGTGGCTTATGCCTATAAT |
| **Chr19F** | TCTCTTCTCTCCCCCACAGA | ATTACAGGCATGCAACACCA |
| | ***Allele-specific Real Time PCR*** | |
| **Exon 7** | CACCCCAGGTTGGCACTG | GACTTGCCCATCTCTGGTTTCC |
| **Probe exon 7 WT** | Fam-AATTGGACGGCGGTTCATCC | |
| **Probe exon 7 R279H** | Hex-AATTGGACGGTGGTTCATCCC | |
| **Exon 8** | GATGAACCGCCGTCCAATTT | CTGTCCGAAACTTGCTGCTT |
| **Probe exon 8 WT** | Fam-CACAGATCCGGGCCTCAAAG | |
| **Probe exon 8 R304Q** | Hex-CACAGATCTGGGCCTCAAAG | |

**Supplementary Table 2:** analysis of TA-cloning on selected edited clones.

| R279H-hiPSCs clone # | N. of wt alleles sequenced and modification | N. of mutant alleles sequenced and modification | Total n. of sequences |
|---|---|---|---|
| **8** | 5 edited<br>3 not edited | 9 edited<br>8 non edited | 25 |
| **11** | 3 edited<br>5 non edited | 1 non edited | 9 |
| **39** | 4 edited | 26 non edited | 30 |
| **54** | 2 edited<br>2 non edited | 4 non edited | 8 |
| **83** | 39 non edited | 9 edited | 48 |
| **156** | none | 8 non edited | 8 |

| R304Q-hiPSCs clone # | N. of wt alleles sequenced and modification | N. of mutant alleles sequenced and modification | Total number of alleles sequences |
|---|---|---|---|
| **10** | 6 non edited | 6 non edited | 12 |
| **42** | | 1 edited<br>11 non edited | 12 |
| **56** | 6 non edited | 3 non edited | 9 |
| **63** | 4 edited | 28 non edited | 32 |
| **73** | 1 edited<br>6 non edited | 2 edited<br>2 non edited | 11 |
| **75** | 1 edited<br>21 non edited | 5 edited<br>1 non edited | 28 |
| **84** | 11 non edited | 1 edited | 12 |

Wt: aattcaacgagggacagattgcccctcctagtcatttgattcgagtagaggggaacagccatgcccagtatgtagaag

**Selected R279H clones**

clone#8: aattcaacgagggacagattgcccctcctaAgtcatttgattcgagtagaggggaacagccatgcccagtatgtagaag +1
clone#11: aattcaacgagggacagatt------------------tgattcgagtagaggggaacagccatgcccagtatgtagaag -14
clone#39: aattcaacgagggacagattgcccctcctaAgtcatttgattcgagtagaggggaacagccatgcccagtatgtagaag +1
clone#54: aattcaacgagggac---------------tagtcatttgattcgagtagaggggaacagccatgcccagtatgtagaag -13
clone#83: aattcaacgagggacag----------------tcatttgattcgagtagaggggaacagccatgcccagtatgtagaag -14
clone#173: aattca acgagggacagatt------------------gattcgagtagagggga acagccatgcccagtatgtagaag -17
clone#156: aattca acgagggacaga-----------tagtcatttgattcgagtagaggggaacagccatgcccagtatgtagaag -10

**Selected R304Q clones**

clone#10 aattcaacg---------------------------catttgattcgagtagaggggaacagccatgcccagtatgtagaag -23
clone#42 aattcaacgagggacagattgcccctcctAagtcatttgattcgagtagaggggaacagccatgcccagtatgtagaag +1
clone#56 aattcaac--------------------------gtcatttgattcgagtagaggggaacagccatgcccagtatgtagaag -22
clone#63 aattcaacgagggacagattgc----------catttgattcgagtagaggggaacagccatgcccagtatgtagaag -10
clone#73 aattcaacgagggacagattgcccctcctAagtcatttgattcgagtagaggggaacagccatgcccagtatgtagaag +1
clone#75 aattcaacgagggacagattgcccctcctAagtcatttgattcgagtagaggggaacagccatgcccagtatgtagaag +1
clone#84 aattcaacgagggacagattgcccctcctAagtcatttgattcgagtagaggggaacagccatgcccagtatgtagaag +1

**Supplementary Figure 1.** Indels-containing sequences of edited R279H-and R304Q-hiPSC selected clones, compared to the wild-type (wt) target region. Numbers on the right represent the number of nucleotides deleted (-) or inserted (+).

```
clustalw.aln

CLUSTAL 2.1 multiple sequence alignment

R279H-hiPSCs-wt+/R279H-            --------------------------------------------------
R304Q-hiPSCs-wt+/R304Q-            --------------------------------------------------
NC_000001.11_217878300-2178795     AGTAAACAAATCATTCTCTCTGAGATTTGTAAATGTGCTGCTCTGCATAG

R279H-hiPSCs-wt+/R279H-            --------------------------------CTTCTATTGGAAGAACTT
R304Q-hiPSCs-wt+/R304Q-            --------------------------------GTTCT-CTGGAG--ACTT
NC_000001.11_217878300-2178795     TTACAATTCAACCAGGTCACCATCACTAGGGCATGAACTTTGAAGTACTT
                                                                    *     * **   ****

R279H-hiPSCs-wt+/R279H-            TTGTTGA-TAAATGTAACAGGTTGTATGTCATCCTTTATCCCCTGATGGC
R304Q-hiPSCs-wt+/R304Q-            TTGTTGA-TAA-TGTAACAGGTTGTATGTCATCCTTTATCCCCTGATGGC
NC_000001.11_217878300-2178795     TTGTTGAATAAATGTAACAGGTTGTATGTCATCCTTTATCCCCTGATGGC
                                   ******* *** **************************************

R279H-hiPSCs-wt+/R279H-            ATCTTTCAGCTACTTAGCTAAGGATGAAGGTGGCATTCTGGTTATGGAAT
R304Q-hiPSCs-wt+/R304Q-            ATCTTTCAGCTACTTAGCTAAGGATGAAGGTGGCATTCTGGTTATGGAAT
NC_000001.11_217878300-2178795     ATCTTTCAGCTACTTAGCTAAGGATGAAGGTGGCATTCTGGTTATGGAAT
                                   **************************************************

R279H-hiPSCs-wt+/R279H-            GAAGTCTGTCTCTGTGTAACACTAACTTGACAGTATAAAAGTAAAGTTTA
R304Q-hiPSCs-wt+/R304Q-            GAAGTCTGTCTCTGTGTAACACTAACTTGACAGTATAAAAGTAAAGTTTA
NC_000001.11_217878300-2178795     GAAGTCTGTCTCTGTGTAACACTAACTTGACAGTATAAAAGTAAAGTTTA
                                   **************************************************

R279H-hiPSCs-wt+/R279H-            TGACATTGGATCACCGATATATTATTATGATCAACTGAGCTAACCAGCCT
R304Q-hiPSCs-wt+/R304Q-            TGACATTGGATCACCGATATATTATTATGATCAACTGAGCTAACCAGCCT
NC_000001.11_217878300-2178795     TGACATTGGATCACCGATATATTATTATGATCAACTGAGCTAACCAGCCT
                                   **************************************************

R279H-hiPSCs-wt+/R279H-            AGACATTTGATTCAAGTTGCATTGTGTATTGCCTGATCGATTCAAAGAAA
R304Q-hiPSCs-wt+/R304Q-            AGACATTTGATTCAAGTTGCATTGTGTATTGCCTGATCGATTCAAAGAAA
NC_000001.11_217878300-2178795     AGACATTTGATTCAAGTTGCATTGTGTATTGCCTGATCGATTCAAAGAAA
                                   **************************************************
```

**Supplementary Figure 2. Absence of off-target editing events on Chromosome 1.** Multiple sequence alignment of edited R279H-hiPSCs$^{wt+/R279H-}$ and R304Q-hiPSCs$^{wt+/R304Q-}$ clones, compared to the predicted off-target site on chromosome 1. Potential off-target editing on chromosome 1 was assessed by amplifying and sequencing the predicted locus identified by the CRISPR design tool (Sigma). The obtained sequences were aligned to the reference human genome using ClustalW software (version 2.1; https://www.genome.jp/tools-bin/clustalw). The boxed sequence indicates the region predicted by the software as a potential off-target site. No sequence mismatches were detected, confirming the absence of off-target editing at this site.

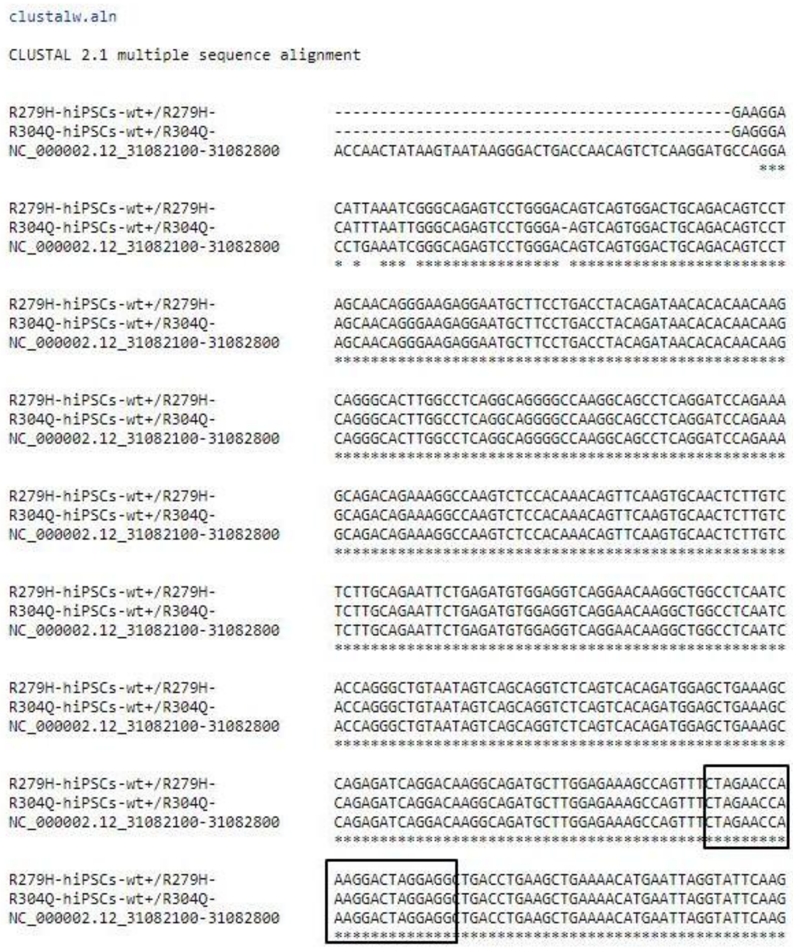

```
clustalw.aln

CLUSTAL 2.1 multiple sequence alignment

R279H-hiPSCs-wt+/R279H-          --------------------------------------------GAAGGA
R304Q-hiPSCs-wt+/R304Q-          --------------------------------------------GAGGGA
NC_000002.12_31082100-31082800   ACCAACTATAAGTAATAAGGGACTGACCAACAGTCTCAAGGATGCCAGGA
                                                                             ***

R279H-hiPSCs-wt+/R279H-          CATTAAATCGGGCAGAGTCCTGGGACAGTCAGTGGACTGCAGACAGTCCT
R304Q-hiPSCs-wt+/R304Q-          CATTTAATTGGGCAGAGTCCTGGGA-AGTCAGTGGACTGCAGACAGTCCT
NC_000002.12_31082100-31082800   CCTGAAATCGGGCAGAGTCCTGGGACAGTCAGTGGACTGCAGACAGTCCT
                                 * *  *** **************** ************************

R279H-hiPSCs-wt+/R279H-          AGCAACAGGGAAGAGGAATGCTTCCTGACCTACAGATAACACACAACAAG
R304Q-hiPSCs-wt+/R304Q-          AGCAACAGGGAAGAGGAATGCTTCCTGACCTACAGATAACACACAACAAG
NC_000002.12_31082100-31082800   AGCAACAGGGAAGAGGAATGCTTCCTGACCTACAGATAACACACAACAAG
                                 **************************************************

R279H-hiPSCs-wt+/R279H-          CAGGGCACTTGGCCTCAGGCAGGGGCCAAGGCAGCCTCAGGATCCAGAAA
R304Q-hiPSCs-wt+/R304Q-          CAGGGCACTTGGCCTCAGGCAGGGGCCAAGGCAGCCTCAGGATCCAGAAA
NC_000002.12_31082100-31082800   CAGGGCACTTGGCCTCAGGCAGGGGCCAAGGCAGCCTCAGGATCCAGAAA
                                 **************************************************

R279H-hiPSCs-wt+/R279H-          GCAGACAGAAAGGCCAAGTCTCCACAAACAGTTCAAGTGCAACTCTTGTC
R304Q-hiPSCs-wt+/R304Q-          GCAGACAGAAAGGCCAAGTCTCCACAAACAGTTCAAGTGCAACTCTTGTC
NC_000002.12_31082100-31082800   GCAGACAGAAAGGCCAAGTCTCCACAAACAGTTCAAGTGCAACTCTTGTC
                                 **************************************************

R279H-hiPSCs-wt+/R279H-          TCTTGCAGAATTCTGAGATGTGGAGGTCAGGAACAAGGCTGGCCTCAATC
R304Q-hiPSCs-wt+/R304Q-          TCTTGCAGAATTCTGAGATGTGGAGGTCAGGAACAAGGCTGGCCTCAATC
NC_000002.12_31082100-31082800   TCTTGCAGAATTCTGAGATGTGGAGGTCAGGAACAAGGCTGGCCTCAATC
                                 **************************************************

R279H-hiPSCs-wt+/R279H-          ACCAGGGCTGTAATAGTCAGCAGGTCTCAGTCACAGATGGAGCTGAAAGC
R304Q-hiPSCs-wt+/R304Q-          ACCAGGGCTGTAATAGTCAGCAGGTCTCAGTCACAGATGGAGCTGAAAGC
NC_000002.12_31082100-31082800   ACCAGGGCTGTAATAGTCAGCAGGTCTCAGTCACAGATGGAGCTGAAAGC
                                 **************************************************

R279H-hiPSCs-wt+/R279H-          CAGAGATCAGGACAAGGCAGATGCTTGGAGAAAGCCAGTTTCTAGAACCA
R304Q-hiPSCs-wt+/R304Q-          CAGAGATCAGGACAAGGCAGATGCTTGGAGAAAGCCAGTTTCTAGAACCA
NC_000002.12_31082100-31082800   CAGAGATCAGGACAAGGCAGATGCTTGGAGAAAGCCAGTTTCTAGAACCA
                                 **************************************************

R279H-hiPSCs-wt+/R279H-          AAGGACTAGGAGGCTGACCTGAAGCTGAAAACATGAATTAGGTATTCAAG
R304Q-hiPSCs-wt+/R304Q-          AAGGACTAGGAGGCTGACCTGAAGCTGAAAACATGAATTAGGTATTCAAG
NC_000002.12_31082100-31082800   AAGGACTAGGAGGCTGACCTGAAGCTGAAAACATGAATTAGGTATTCAAG
                                 **************************************************
```

**Supplementary Figure 3. Absence of off-target editing events on Chromosome 2.** Multiple sequence alignment of edited R279H-hiPSC$^{wt+/R279H-}$ and R304Q-hiPSC$^{wt+/R304Q-}$ clones, compared to the predicted off-target site on chromosome 2. Potential off-target editing on chromosome 2 was assessed by amplifying and sequencing the predicted locus identified by the CRISPR design tool (Sigma). The obtained sequences were aligned to the reference human genome using ClustalW software (version 2.1; https://www.genome.jp/tools-bin/clustalw). The boxed sequence indicates the region predicted by the software as a potential off-target site. No sequence mismatches were detected, confirming the absence of off-target editing at this site.

```
clustalw.aln

CLUSTAL 2.1 multiple sequence alignment

R279H-hiPSCs-wt+/R279H-            --------------------------------------------------
R304Q-hiPSCs-wt+/R304Q-            --------------------------------------------------
NC_000003.12_193439500-1934404     ATGGGAAAGTACTTATAAAGTCTAAAATGATAACAAGTCATAGTTGTTTG

R279H-hiPSCs-wt+/R279H-            --------------------------------------------------
R304Q-hiPSCs-wt+/R304Q-            --------------------------------------------------
NC_000003.12_193439500-1934404     TGCTGTTAATACTAGCATGAGAGGAAATCCATCCTGTCAACAGTCACTCA

R279H-hiPSCs-wt+/R279H-            ---------------------------------------------GCTCG
R304Q-hiPSCs-wt+/R304Q-            --------------------------------------------AAGTCG
NC_000003.12_193439500-1934404     ACTCAAGTATGTAATGCCAGGCCTAGAAGTTGAAGAAAGAAAACACATCC
                                                                                  ** 

R279H-hiPSCs-wt+/R279H-            TTGCTTTAGA-TGACTTTTTGAGGCTGGGTGAATACTCTCTTGAATCATT
R304Q-hiPSCs-wt+/R304Q-            TTGCTTTAGA-TGACTTTTTGAGGCTGGGTGAATACTCTCTTGAATCATT
NC_000003.12_193439500-1934404     TTGTTTCAGAATGACTTTTTGAGGCTGGGTGAATACTCTCTTGAATCATT
                                   *** ** *** ***************************************

R279H-hiPSCs-wt+/R279H-            GACAAGGACTATTGAGAGCCAACTTTTTAAATTATGAGCATTCTGACAGC
R304Q-hiPSCs-wt+/R304Q-            GACAAGGACTATTGAGAGCCAACTTTTTAAATTATGAGCATTCTGACAGC
NC_000003.12_193439500-1934404     GACAAGGACTATTGAGAGCCAACTTTTTAAATTATGAGCATTCTGACAGC
                                   **************************************************

R279H-hiPSCs-wt+/R279H-            ATTCATAAAACATGCTATACTTTTCTTCCACACTGCCATATGGAGGGAAC
R304Q-hiPSCs-wt+/R304Q-            ATTCATAAAACATGCTATACTTTTCTTCCACACTGCCATATGGAGGGAAC
NC_000003.12_193439500-1934404     ATTCATAAAACATGCTATACTTTTCTTCCACACTGCCATATGGAGGGAAC
                                   **************************************************

R279H-hiPSCs-wt+/R279H-            TCGTTGACTCTCTAGGTTTCAGCCATGTTGAGAAGATAGGGCTTTCTGAG
R304Q-hiPSCs-wt+/R304Q-            TCGTTGACTCTCTAGGTTTCAGCCATGTTGAGAAGATAGGGCTTTCTGAG
NC_000003.12_193439500-1934404     TCGTTGACTCTCTAGGTTTCAGCCATGTTGAGAAGATAGGGCTTTCTGAG
                                   **************************************************

R279H-hiPSCs-wt+/R279H-            CTGAGAAGCTAGCCTGATAACGGAATCAAATGAATAGGAGGGCAGAGCAG
R304Q-hiPSCs-wt+/R304Q-            CTGAGAAGCTAGCCTGATAACGGAATCAAATGAATAGGAGGGCAGAGCAG
NC_000003.12_193439500-1934404     CTGAGAAGCTAGCCTGATAACGGAATCAAATGAATAGGAGGGCAGAGCAG
                                   **************************************************
```

**Supplementary Figure 4. Absence of off-target editing events on Chromosome 3.** Multiple sequence alignment of edited R279H-hiPSC$^{wt+/R279H-}$ and R304Q-hiPSC$^{wt+/R304Q-}$ clones, compared to the predicted off-target site on chromosome 3. Potential off-target editing on chromosome 3 was assessed by amplifying and sequencing the predicted locus identified by the CRISPR design tool (Sigma). The obtained sequences were aligned to the reference human genome using ClustalW software (version 2.1; https://www.genome.jp/tools-bin/clustalw). The boxed sequence indicates the region predicted by the software as a potential off-target site. No sequence mismatches were detected, confirming the absence of off-target editing at this site.

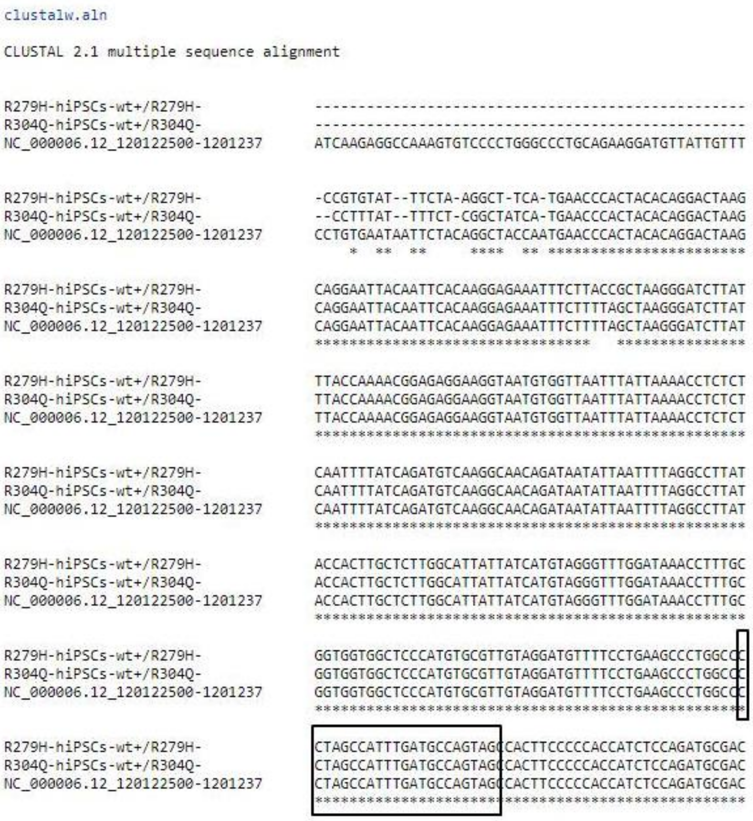

```
clustalw.aln

CLUSTAL 2.1 multiple sequence alignment

R279H-hiPSCs-wt+/R279H-             --------------------------------------------------
R304Q-hiPSCs-wt+/R304Q-             --------------------------------------------------
NC_000006.12_120122500-1201237      ATCAAGAGGCCAAAGTGTCCCCTGGGCCCTGCAGAAGGATGTTATTGTTT

R279H-hiPSCs-wt+/R279H-             -CCGTGTAT--TTCTA-AGGCT-TCA-TGAACCCACTACACAGGACTAAG
R304Q-hiPSCs-wt+/R304Q-             --CCTTTAT--TTTCT-CGGCTATCA-TGAACCCACTACACAGGACTAAG
NC_000006.12_120122500-1201237      CCTGTGAATAATTCTACAGGCTACCAATGAACCCACTACACAGGACTAAG
                                        *  **  **     ****  ** ***********************

R279H-hiPSCs-wt+/R279H-             CAGGAATTACAATTCACAAGGAGAAATTTCTTACCGCTAAGGGATCTTAT
R304Q-hiPSCs-wt+/R304Q-             CAGGAATTACAATTCACAAGGAGAAATTTCTTTAGCTAAGGGATCTTAT
NC_000006.12_120122500-1201237      CAGGAATTACAATTCACAAGGAGAAATTTCTTTAGCTAAGGGATCTTAT
                                    ********************************   ***************

R279H-hiPSCs-wt+/R279H-             TTACCAAAACGGAGAGGAAGGTAATGTGGTTAATTTATTAAAACCTCTCT
R304Q-hiPSCs-wt+/R304Q-             TTACCAAAACGGAGAGGAAGGTAATGTGGTTAATTTATTAAAACCTCTCT
NC_000006.12_120122500-1201237      TTACCAAAACGGAGAGGAAGGTAATGTGGTTAATTTATTAAAACCTCTCT
                                    **************************************************

R279H-hiPSCs-wt+/R279H-             CAATTTTATCAGATGTCAAGGCAACAGATAATATTAATTTTAGGCCTTAT
R304Q-hiPSCs-wt+/R304Q-             CAATTTTATCAGATGTCAAGGCAACAGATAATATTAATTTTAGGCCTTAT
NC_000006.12_120122500-1201237      CAATTTTATCAGATGTCAAGGCAACAGATAATATTAATTTTAGGCCTTAT
                                    **************************************************

R279H-hiPSCs-wt+/R279H-             ACCACTTGCTCTTGGCATTATTATCATGTAGGGTTTGGATAAACCTTTGC
R304Q-hiPSCs-wt+/R304Q-             ACCACTTGCTCTTGGCATTATTATCATGTAGGGTTTGGATAAACCTTTGC
NC_000006.12_120122500-1201237      ACCACTTGCTCTTGGCATTATTATCATGTAGGGTTTGGATAAACCTTTGC
                                    **************************************************

R279H-hiPSCs-wt+/R279H-             GGTGGTGGCTCCCATGTGCGTTGTAGGATGTTTTCCTGAAGCCCTGGCCC
R304Q-hiPSCs-wt+/R304Q-             GGTGGTGGCTCCCATGTGCGTTGTAGGATGTTTTCCTGAAGCCCTGGCCC
NC_000006.12_120122500-1201237      GGTGGTGGCTCCCATGTGCGTTGTAGGATGTTTTCCTGAAGCCCTGGCCC
                                    **************************************************

R279H-hiPSCs-wt+/R279H-             CTAGCCATTTGATGCCAGTAGCCACTTCCCCCACCATCTCCAGATGCGAC
R304Q-hiPSCs-wt+/R304Q-             CTAGCCATTTGATGCCAGTAGCCACTTCCCCCACCATCTCCAGATGCGAC
NC_000006.12_120122500-1201237      CTAGCCATTTGATGCCAGTAGCCACTTCCCCCACCATCTCCAGATGCGAC
                                    **************************************************
```

**Supplementary Figure 5. Absence of off-target editing events on Chromosome 6.** Multiple sequence alignment of edited R279H-hiPSC$^{wt+/R279H-}$ and R304Q-hiPSC$^{wt+/R304Q-}$ clones, compared to the predicted off-target site on chromosome 6. Potential off-target editing on chromosome 6 was assessed by amplifying and sequencing the predicted locus identified by the CRISPR design tool (Sigma). The obtained sequences were aligned to the reference human genome using ClustalW software (version 2.1; https://www.genome.jp/tools-bin/clustalw). The boxed sequence indicates the region predicted by the software as a potential off-target site. No sequence mismatches were detected, confirming the absence of off-target editing at this site.

```
clustalw.aln

CLUSTAL 2.1 multiple sequence alignment

R279H-hiPSCs-wt+/R279H-         ----------------------------------------------GGGA
R304Q-hiPSCs-wt+/R304Q-         ----------------------------------------------CGGA
NC_000008.11_74801300-74801900  GAACCATGTGTATAAATCGGGATTCTACAGAGAAACAGAATCAATACAAG

R279H-hiPSCs-wt+/R279H-         ATGGAAATAGGGATAGGAGAGATAGATAGATAGATTGATTGATTGATTTG
R304Q-hiPSCs-wt+/R304Q-         ATGGATATAGATA---GAGAGATAGATAGATAGATTGATTGATTGATTTG
NC_000008.11_74801300-74801900  ATAGATATAGATA---GAGAGATAGATAGATAGATTGATTGATTGATTTG
                                ** ** ****  *   **********************************

R279H-hiPSCs-wt+/R279H-         TTTTCAGTAATTGGCTCATGCAATTATAGAAGCTGAAAAGTCCCAAGATC
R304Q-hiPSCs-wt+/R304Q-         TTTTCAGTAATTGGCTCATGCAATTATAGAAGCTGAAAAGTCCCAAGATC
NC_000008.11_74801300-74801900  TTTTCAGTAATTGGCTCATGCAATTATAGAAGCTGAAAAGTCCCAAGATC
                                **************************************************

R279H-hiPSCs-wt+/R279H-         TGTAACTGGCAAGCTGGAGACCCAGGAGAGCTTGTTGTGTAGTTCCAGTT
R304Q-hiPSCs-wt+/R304Q-         TGTAACTGGCAAGCTGGAGACCCAGGAGAGCTTGTTGTGTAGTTCCAGTT
NC_000008.11_74801300-74801900  TGTAACTGGCAAGCTGGAGACCCAGGAGAGCTTGTTGTGTAGTTCCAGTT
                                **************************************************

R279H-hiPSCs-wt+/R279H-         TGCAAGCTGGTAGGCTCAAGACCAAGGAAGGGTAGATGTTTCAGTTTGAG
R304Q-hiPSCs-wt+/R304Q-         TGCAAGCTGGTAGGCTCAAGACCAAGGAAGGGTAGATGTTTCAGTTTGAG
NC_000008.11_74801300-74801900  TGCAAGCTGGTAGGCTCAAGACCAAGGAAGGGTAGATGTTTCAGTTTGAG
                                **************************************************

R279H-hiPSCs-wt+/R279H-         TCCGAAGGCAGAATAAAGACCAAAGTCCCAGCTCAAGCAGTCAGGCAGGA
R304Q-hiPSCs-wt+/R304Q-         TCCGAAGGCAGAATAAAGACCAAAGTCCCAGCTCAAGCAGTCAGGCAGGA
NC_000008.11_74801300-74801900  TCCGAAGGCAGAATAAAGACCAAAGTCCCAGCTCAAGCAGTCAGGCAGGA
                                **************************************************

R279H-hiPSCs-wt+/R279H-         GGAAGACCCTCTTAGTCATTTTATTCTATTTAGGTCTTCAGCTGGATGGA
R304Q-hiPSCs-wt+/R304Q-         GGAAGACCCTCTTAGTCATTTTATTCTATTTAGGTCTTCAGCTGGATGGA
NC_000008.11_74801300-74801900  GGAAGACCCTCTTAGTCATTTTATTCTATTTAGGTCTTCAGCTGGATGGA
                                **************************************************

R279H-hiPSCs-wt+/R279H-         TGAGGCCCATCTATAGTAGGGAGGACAATCTACTTTAATCAGTCTACTGA
R304Q-hiPSCs-wt+/R304Q-         TGAGGCCCATCTATAGTAGGGAGGACAATCTACTTTAATCAGTCTACTGA
NC_000008.11_74801300-74801900  TGAGGCCCATCTATAGTAGGGAGGACAATCTACTTTAATCAGTCTACTGA
                                **************************************************
```

**Supplementary Figure 6. Absence of off-target editing events on Chromosome 8.** Multiple sequence alignment of edited R279H-hiPSC$^{wt+/R279H-}$ and R304Q-hiPSC$^{wt+/R304Q-}$ clones, compared to the predicted off-target site on chromosome 8. Potential off-target editing on chromosome 8 was assessed by amplifying and sequencing the predicted locus identified by the CRISPR design tool (Sigma). The obtained sequences were aligned to the reference human genome using ClustalW software (version 2.1; https://www.genome.jp/tools-bin/clustalw). The boxed sequence indicates the region predicted by the software as a potential off-target site. No sequence mismatches were detected, confirming the absence of off-target editing at this site.

```
clustalw.aln

CLUSTAL 2.1 multiple sequence alignment

NC_000010.11_126175300-1261759      GGAGTTGCAACTCACCTCCACAATGGCCTGCCTGCTGGCGGGGCAACCTT
R279H-hiPSCs-wt+/R279H-             --------------------------------------------------
R304Q-hiPSCs-wt+/R304Q-             --------------------------------------------------

NC_000010.11_126175300-1261759      ACAAAATATTATTATTATTTACTCATAACAATAACAATTGCTCGGAAAAT
R279H-hiPSCs-wt+/R279H-             -----------------TTTACTTATA--CATAACAATTGCTCGGAAA-T
R304Q-hiPSCs-wt+/R304Q-             ------------------TTTCTTTTA--AATAACAATTGCTCGGAAAAT
                                                      ** **  **   ****************** *

NC_000010.11_126175300-1261759      AAAACTCAAAACAGGCAAAACCATTGGAACACCAAGACGAGACTTCTACA
R279H-hiPSCs-wt+/R279H-             AAAACTCAAAACAGGCAAAACCATTGGAACACCAAGACGAGACTTCTACA
R304Q-hiPSCs-wt+/R304Q-             AAAACTCAAAACAGGCAAAACCATTGGAACACCAAGACGAGACTTCTACA
                                    **************************************************

NC_000010.11_126175300-1261759      GGTTTTATTTTAAACATATGTTTCTAAAACAGTCAGTGGTGAGTGCCAAA
R279H-hiPSCs-wt+/R279H-             GGTTTTATTTTAAACATATGTTTCTAAAACAGTCAGTGGTGAGTGCCAAA
R304Q-hiPSCs-wt+/R304Q-             GGTTTTATTTTAAACATATGTTTCTAAAACAGTCAGTGGTGAGTGCCAAA
                                    **************************************************

NC_000010.11_126175300-1261759      AAACACAGCATCAAGTCGATTTTCCTTTTTCTGTGTACATGACTTGCTTT
R279H-hiPSCs-wt+/R279H-             AAACACAGCATCAAGTCGATTTTCCTTTTTCTGTGTACATGACTTGCTTT
R304Q-hiPSCs-wt+/R304Q-             AAACACAGCATCAAGTCGATTTTCCTTTTTCTGTGTACATGACTTGCTTT
                                    **************************************************

NC_000010.11_126175300-1261759      GGAAACTGCCTATACATGCCCTACATTTTTTTCTCTCATCAAAAACAATC
R279H-hiPSCs-wt+/R279H-             GGAAACTGCCTATACATGCCCTACATTTTTTTCTCTCATCAAAAACAATC
R304Q-hiPSCs-wt+/R304Q-             GGAAACTGCCTATACATGCCCTACATTTTTTTCTCTCATCAAAAACAATC
                                    **************************************************

NC_000010.11_126175300-1261759      GAGGAATGGTACAAGGAGTTCTGAAGCATAACCTAGTCGTTTCATTCGAG
R279H-hiPSCs-wt+/R279H-             GAGGAATGGTACAAGGAGTTCTGAAGCATAACCTAGTCGTTTCATTCGAG
R304Q-hiPSCs-wt+/R304Q-             GAGGAATGGTACAAGGAGTTCTGAAGCATAACCTAGTCGTTTCATTCGAG
                                    **************************************************

NC_000010.11_126175300-1261759      TGGCGAGAGAGGGACGTGATGTGATGTGGGTGCCACGTAAGTGTCGTGCT
R279H-hiPSCs-wt+/R279H-             TGGCGAGAGAGGGACGTGATGTGATGTGGGTGCCACGTAAGTGTCGTGCT
R304Q-hiPSCs-wt+/R304Q-             TGGCGAGAGAGGGACGTGATGTGATGTGGGTGCCACGTAAGTGTCGTGCT
                                    **************************************************
```

**Supplementary Figure 7. Absence of off-target editing events on Chromosome 10.** Multiple sequence alignment of edited R279H-hiPSC$^{wt+/R279H-}$ and R304Q-hiPSC$^{wt+/R304Q-}$ clones, compared to the predicted off-target site on chromosome 10. Potential off-target editing on chromosome 10 was assessed by amplifying and sequencing the predicted locus identified by the CRISPR design tool (Sigma). The obtained sequences were aligned to the reference human genome using ClustalW software (version 2.1; https://www.genome.jp/tools-bin/clustalw). The boxed sequence indicates the region predicted by the software as a potential off-target site. No sequence mismatches were detected, confirming the absence of off-target editing at this site.

```
clustalw.aln

CLUSTAL 2.1 multiple sequence alignment

NC_000012.12_64737000-64737800      CTACCTTGCTTGTCTGATTTGGTTTGATCCCAGGTCCTCGAACCAACAGT
R304Q-hiPSCs-wt+/R304Q-             --------------------------------------------------
R279H-hiPSCs-wt+/R279H-             --------------------------------------------------

NC_000012.12_64737000-64737800      GGAACTTTGATATCAAACTCATACAGCTGTCTCTTGTCTATTGGCAAGGA
R304Q-hiPSCs-wt+/R304Q-             --------------------------------------------------
R279H-hiPSCs-wt+/R279H-             --------------------------------------------------

NC_000012.12_64737000-64737800      AAACTGTCCTGAAAACAAAACACACAATG-TAAAGATGGAGCCAAGACAG
R304Q-hiPSCs-wt+/R304Q-             -----------------------GAAATGCTAAAGATGGAGCCAAGACAG
R279H-hiPSCs-wt+/R279H-             ------------------------CAACG-CTAAGATGGAGCCA-GACAG
                                                             ** *   ************ *****

NC_000012.12_64737000-64737800      GAGCCTTGCTCATGTTATGTTTCACTCATTTAATCCACAGCCAAAACTAA
R304Q-hiPSCs-wt+/R304Q-             GAGCCTTGCTCATGTTATGTTTCACTCATTTAATCCACAGCCAAAACTAA
R279H-hiPSCs-wt+/R279H-             GAGCCTTGCTCATGTTATGTTTCACTCATTTAATCCACAGCCAAAACTAA
                                    **************************************************

NC_000012.12_64737000-64737800      AACAACTTTGTTCTACTGATTCTGGTGTTTTAATAGAAAGCTGAAATTTG
R304Q-hiPSCs-wt+/R304Q-             AACAACTTTGTTCTACTGATTCTGGTGTTTTAATAGAAAGCTGAAATTTG
R279H-hiPSCs-wt+/R279H-             AACAACTTTGTTCTACTGATTCTGGTGTTTTAATAGAAAGCTGAAATTTG
                                    **************************************************

NC_000012.12_64737000-64737800      CCTGGGTCTCTTGTTACTACAAGGCAGGAAAACAACAAATTCATCTTCTC
R304Q-hiPSCs-wt+/R304Q-             CCTGGGTCTCTTGTTACTACAAGGCAGGAAAACAACAAATTCATCTTCTC
R279H-hiPSCs-wt+/R279H-             CCTGGGTCTCTTGTTACTACAAGGCAGGAAAACAACAAATTCATCTTCTC
                                    **************************************************

NC_000012.12_64737000-64737800      TAAACAAATGACTAGAAGGGACTCCCTGGCTTATGATAGAACAAAATAAG
R304Q-hiPSCs-wt+/R304Q-             TAAACAAATGACTAGAAGGGACTCCCTGGCTTATGATAGAACAAAATAAG
R279H-hiPSCs-wt+/R279H-             TAAACAAATGACTAGAAGGGACTCCCTGGCTTATGATAGAACAAAATAAG
                                    **************************************************

NC_000012.12_64737000-64737800      CAAAAATAATGACAGGAAAGTATTGAAACACAAATACATAAAAAATTATG
R304Q-hiPSCs-wt+/R304Q-             CAAAAATAATGACAGGAAAGTATTGAAACACAAATACATAAAAAATTATG
R279H-hiPSCs-wt+/R279H-             CAAAAATAATGACAGGAAAGTATTGAAACACAAATACATAAAAAATTATG
                                    **************************************************
```

**Supplementary Figure 8. Absence of off-target editing events on Chromosome 12.** Multiple sequence alignment of edited R279H-hiPSC$^{wt+/R279H-}$ and R304Q-hiPSC$^{wt+/R304Q-}$ clones, compared to the predicted off-target site on chromosome 12. Potential off-target editing on chromosome 12 was assessed by amplifying and sequencing the predicted locus identified by the CRISPR design tool (Sigma). The obtained sequences were aligned to the reference human genome using ClustalW software (version 2.1; https://www.genome.jp/tools-bin/clustalw). The boxed sequence indicates the region predicted by the software as a potential off-target site. No sequence mismatches were detected, confirming the absence of off-target editing at this site.

```
clustalw.aln

CLUSTAL 2.1 multiple sequence alignment

NC_000014.9_39883300-39883900      ACAATACCCATTTGGGCTTCTTTTCATTAACACATCCATATAACTTAAAA
R304Q-hiPSCs-wt+/R304Q-            ------------------------------------CCTAAAACTTAAA-
R279H-hiPSCs-wt+/R279H-            ----------------------------------CTTTTTAACCTTTAAA
                                                                       **   ** **

NC_000014.9_39883300-39883900      GTATATGTTATTACTGATTTTCTTGGCCAGATATTTCTCCATCCCTCCTC
R304Q-hiPSCs-wt+/R304Q-            GTATATGTTATTACTGATTTTCTTGGCCAGATATTTCTCCATCCCTCCTC
R279H-hiPSCs-wt+/R279H-            GTATATGTTATTACTGATTTTCTTGGCCAGATATTTCTCCATCCCTCCTC
                                   **************************************************

NC_000014.9_39883300-39883900      TCTTACCACCCACTCATATACATCGGAACAACTTTTTATTATAGTTCATC
R304Q-hiPSCs-wt+/R304Q-            TCTTACCACCCACTCATATACATCGGAACAACTTTTTATTATAGTTCATC
R279H-hiPSCs-wt+/R279H-            TCTTACCACCCACTCATATACATCGGAACAACTTTTTATTATAGTTCATC
                                   **************************************************

NC_000014.9_39883300-39883900      TCAAATTAAGGAAAACATAGAGTATCCCTTGTGGAGGTAATGAATTACCT
R304Q-hiPSCs-wt+/R304Q-            TCAAATTAAGGAAAACATAGAGTATCCCTTGTGGAGGTAATGAATTACCT
R279H-hiPSCs-wt+/R279H-            TCAAATTAAGGAAAACATAGAGTATCCCTTGTGGAGGTAATGAATTACCT
                                   **************************************************

NC_000014.9_39883300-39883900      GTTAACTATGGTCAAGTTCAGACTAAATGACAACTTTGACATTGCAGAGG
R304Q-hiPSCs-wt+/R304Q-            GTTAACTATGGTCAAGTTCAGACTAAATGACAACTTTGACATTGCAGAGG
R279H-hiPSCs-wt+/R279H-            GTTAACTATGGTCAAGTTCAGACTAAATGACAACTTTGACATTGCAGAGG
                                   **************************************************

NC_000014.9_39883300-39883900      AGATTTAAAATTCAAGGTTAACCTGGGTTACTGGATCAAATCACCTGTCC
R304Q-hiPSCs-wt+/R304Q-            AGATTTAAAATTCAAGGTTAACCTGGGTTACTGGATCAAATCACCTGTCC
R279H-hiPSCs-wt+/R279H-            AGATTTAAAATTCAAGGTTAACCTGGGTTACTGGATCAAATCACCTGTCC
                                   **************************************************

NC_000014.9_39883300-39883900      CTCCTAATCATTTCATTCTAAATGTGCATGAATTATGGTCAGAGTATCT
R304Q-hiPSCs-wt+/R304Q-            CTCCTAATCATTTCATTCTAAATGTGCATGAATTATGGTCAGAGTATCT
R279H-hiPSCs-wt+/R279H-            CTCCTAATCATTTCATTCTAAATGTGCATGAATTATGGTCAGAGTATCT
                                   **************************************************

NC_000014.9_39883300-39883900      CTTCTATTGCCTATTCTAACCCTGATATTTCACACCTTAAGCTGGGGATG
R304Q-hiPSCs-wt+/R304Q-            CTTCTATTGCCTATTCTAACCCTGATATTTCACACCTTAAGCTGGGGATG
R279H-hiPSCs-wt+/R279H-            CTTCTATTGCCTATTCTAACCCTGATATTTCACACCTTAAGCTGGGGATG
                                   **************************************************
```

**Supplementary Figure 9. Absence of off-target editing events on Chromosome 14.** Multiple sequence alignment of edited R279H-hiPSC$^{wt+/R279H-}$ and R304Q-hiPSC$^{wt+/R304Q-}$ clones, compared to the predicted off-target site on chromosome 14. Potential off-target editing on chromosome 14 was assessed by amplifying and sequencing the predicted locus identified by the CRISPR design tool (Sigma). The obtained sequences were aligned to the reference human genome using ClustalW software (version 2.1; https://www.genome.jp/tools-bin/clustalw). The boxed sequence indicates the region predicted by the software as a potential off-target site. No sequence mismatches were detected, confirming the absence of off-target editing at this site.

```
clustalw.aln

CLUSTAL 2.1 multiple sequence alignment

R279H-hiPSCs-wt+/R279H-             --------------------------------------------------
R304Q-hiPSCs-wt+/R304Q-             --------------------------------------------------
NC_000017.11_43516000-43516800      GTTTTTGAAGTAGTGTTTTATTATTTTTGAAAAACAGAAGAAAAAAATTT

R279H-hiPSCs-wt+/R279H-             ------------------------GCTTAAATTTTGCTGTTTTTCATTCC
R304Q-hiPSCs-wt+/R304Q-             --------------------------CTAAATTTT-GCTGTTTTA---TTC
NC_000017.11_43516000-43516800      ACCCACCATCCTACTACCTCCACTGTTATTATTTTGCTGTTTTTCATTCC
                                                              **   *** ********    * *

R279H-hiPSCs-wt+/R279H-             AGTCTTTTTTCATGTTTATTTCTTTCATGGTTGCAATTTTAGTAGTCAAG
R304Q-hiPSCs-wt+/R304Q-             AGTCTTTTTTCATGTTTATTTCTTTCATGGTTGCAATTTTAGTAGTCAAG
NC_000017.11_43516000-43516800      AGTCTTTTTTCATGTTTATTTCTTTCATGGTTGCAATTTTAGTAGTCAAG
                                    **************************************************

R279H-hiPSCs-wt+/R279H-             TAAGTATGCAGTTTTGTATATTTTTTTGAGTATTTAATTTTCCTATTAGA
R304Q-hiPSCs-wt+/R304Q-             TAAGTATGCAGTTTTGTATATTTTTTTGAGTATTTAATTTTCCTATTAGA
NC_000017.11_43516000-43516800      TAAGTATGCAGTTTTGTATATTTTTTTGAGTATTTAATTTTCCTATTAGA
                                    **************************************************

R279H-hiPSCs-wt+/R279H-             ATAATACATGTTCAATATTTGGTAAAATCTAAAGAAAATAGACCTCATCT
R304Q-hiPSCs-wt+/R304Q-             ATAATACATGTTCAATATTTGGTAAAATCTAAAGAAAATAGACCTCATCT
NC_000017.11_43516000-43516800      ATAATACATGTTCAATATTTGGTAAAATCTAAAGAAAATAGACCTCATCT
                                    **************************************************

R279H-hiPSCs-wt+/R279H-             GCAAGCCTACTATCAGAGTTAGCCACTGTTAACATTTTAGGGTTTTTTTC
R304Q-hiPSCs-wt+/R304Q-             GCAAGCCTACTATCAGAGTTAGCCACTGTTAACATTTTAGGGTTTTTTTC
NC_000017.11_43516000-43516800      GCAAGCCTACTATCAGAGTTAGCCACTGTTAACATTTTAGGGTTTTTTTC
                                    **************************************************

R279H-hiPSCs-wt+/R279H-             GCCTAGTCATTTCCTTCTAGTAGTTGATATATAAAGACTTTTACTAAAAT
R304Q-hiPSCs-wt+/R304Q-             GCCTAGTCATTTCCTTCTAGTAGTTGATATATAAAGACTTTTACTAAAAT
NC_000017.11_43516000-43516800      GCCTAGTCATTTCCTTCTAGTAGTTGATATATAAAGACTTTTACTAAAAT
                                    **************************************************

R279H-hiPSCs-wt+/R279H-             TATTTTACATTGTTTCGTAACTTGCATTTTGCATTTAATGCGAATGAACA
R304Q-hiPSCs-wt+/R304Q-             TATTTTACATTGTTTCGTAACTTGCATTTTGCATTTAATGCGAATGAACA
NC_000017.11_43516000-43516800      TATTTTACATTGTTTCGTAACTTGCATTTTGCATTTAATGCGAATGAACA
                                    **************************************************
```

**Supplementary Figure 10. Absence of off-target editing events on Chromosome 17.** Multiple sequence alignment of edited R279H-hiPSC$^{wt+/R279H-}$ and R304Q-hiPSC$^{wt+/R304Q-}$ clones, compared to the predicted off-target site on chromosome 17. Potential off-target editing on chromosome 17 was assessed by amplifying and sequencing the predicted locus identified by the CRISPR design tool (Sigma). The obtained sequences were aligned to the reference human genome using ClustalW software (version 2.1; https://www.genome.jp/tools-bin/clustalw). The boxed sequence indicates the region predicted by the software as a potential off-target site. No sequence mismatches were detected, confirming the absence of off-target editing at this site.

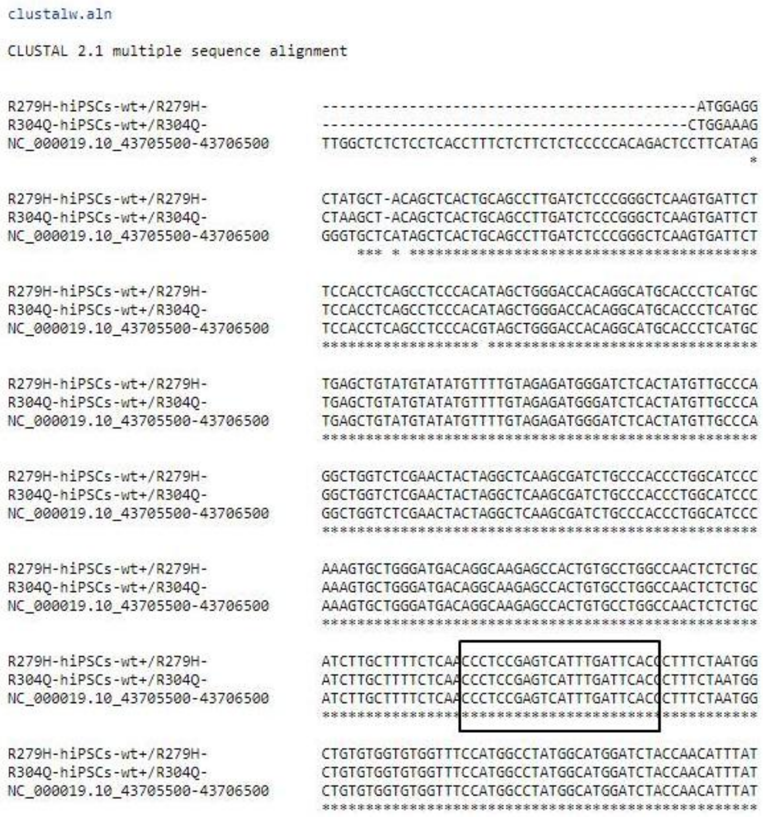


**Supplementary Figure 11. Absence of off-target editing events on Chromosome 19.** Multiple sequence alignment of edited R279H-hiPSC$^{wt+/R279H-}$ and R304Q-hiPSC$^{wt+/R304Q-}$ clones, compared to the predicted off-target site on chromosome 19. Potential off-target editing on chromosome 19 was assessed by amplifying and sequencing the predicted locus identified by the CRISPR design tool (Sigma). The obtained sequences were aligned to the reference human genome using ClustalW software (version 2.1; https://www.genome.jp/tools-bin/clustalw). The boxed sequence indicates the region predicted by the software as a potential off-target site. No sequence mismatches were detected, confirming the absence of off-target editing at this site.

```
clustalw.aln

CLUSTAL 2.1 multiple sequence alignment

NC_000023.11_45315300-45315999      TCCATATATAACAACTATGCATCATATAAAATAAAGATCATATTCTTTTT
R304Q-hiPSCs-wt+/R304Q-             ---------CATAAACTTGCATCATATAAAATAAAGATCATATTCCTTTT
R279H-hiPSCs-wt+/R279H-             --------CTTTCACAATGCATCATATAAAATAAAGATCATATTCCTTTT
                                                 *   **************************** ****

NC_000023.11_45315300-45315999      CCTTCAAATATGTCTTCTAGTATTTTACATTGCAGAAAAGGATTCTAAAC
R304Q-hiPSCs-wt+/R304Q-             CCTTCAAATATGTCTTCTAGTATTTTACATTGCAGAAAAGGATTCTAAAC
R279H-hiPSCs-wt+/R279H-             CCTTCAAATATGTCTTCTAGTATTTTACATTGCAGAAAAGGATTCTAAAC
                                    **************************************************

NC_000023.11_45315300-45315999      CCAGCTTCGTTTTTGTTCTTTTTTCTTTTGTAATTTGTTTTTCATGTCTG
R304Q-hiPSCs-wt+/R304Q-             CCAGCTTCGTTTTTGTTCTTTTTTCTTTTGTAATTTGTTTTTCATGTCTG
R279H-hiPSCs-wt+/R279H-             CCAGCTTCGTTTTTGTTCTTTTTTCTTTTGTAATTTGTTTTTCATGTCTG
                                    **************************************************

NC_000023.11_45315300-45315999      GGTGTTTGTACAATTTCCTTTCCCCTTACAATGAAAAACAGAATGACCAA
R304Q-hiPSCs-wt+/R304Q-             GGTGTTTGTACAATTTCCTTTCCCCTTACAATGAAAAACAGAATGACCAA
R279H-hiPSCs-wt+/R279H-             GGTGTTTGTACAATTTCCTTTCCCCTTACAATGAAAAACAGAATGACCAA
                                    **************************************************

NC_000023.11_45315300-45315999      GATTGATATTTCTAGAGATGTAGTGAGCTCTCTATACCTGTAAATGTAGG
R304Q-hiPSCs-wt+/R304Q-             GATTGATATTTCTAGAGATGTAGTGAGCTCTCTATACCTGTAAATGTAGG
R279H-hiPSCs-wt+/R279H-             GATTGATATTTCTAGAGATGTAGTGAGCTCTCTATACCTGTAAATGTAGG
                                    **************************************************

NC_000023.11_45315300-45315999      GAATTAACTTTCATTGGGTGCCTCTTTAACATCCACTTCTTTCTCTCCTC
R304Q-hiPSCs-wt+/R304Q-             GAATTAACTTTCATTGGGTGCCTCTTTAACATCCACTTCTTTCTCTCCTC
R279H-hiPSCs-wt+/R279H-             GAATTAACTTTCATTGGGTGCCTCTTTAACATCCACTTCTTTCTCTCCTC
                                    **************************************************

NC_000023.11_45315300-45315999      TTCTTAAAATGACTCCACATGTCCTAGCTATTTGATTTGAGTAGATTATT
R304Q-hiPSCs-wt+/R304Q-             TTCTTAAAATGACTCCACATGTCCTAGCTATTTGATTTGAGTAGATTATT
R279H-hiPSCs-wt+/R279H-             TTCTTAAAATGACTCCACATGTCCTAGCTATTTGATTTGAGTAGATTATT
                                    **************************************************

NC_000023.11_45315300-45315999      GATATGAGACCTTTCTTCTTTTTAAAGATAAACATTTAATATTCTAAATT
R304Q-hiPSCs-wt+/R304Q-             GATATGAGACCTTTCTTCTTTTTAAAGATAAACATTTAATATTCTAAATT
R279H-hiPSCs-wt+/R279H-             GATATGAGACCTTTCTTCTTTTTAAAGATAAACATTTAATATTCTAAATT
                                    **************************************************
```

**Supplementary Figure 12. Absence of off-target editing events on Chromosome X.** Multiple sequence alignment of edited R279H-hiPSC$^{wt+/R279H-}$ and R304Q-hiPSC$^{wt+/R304Q-}$ clones, compared to the predicted off-target site on chromosome X. Potential off-target editing on chromosome X was assessed by amplifying and sequencing the predicted locus identified by the CRISPR design tool (Sigma). The obtained sequences were aligned to the reference human genome using ClustalW software (version 2.1; https://www.genome.jp/tools-bin/clustalw). The boxed sequence indicates the region predicted by the software as a potential off-target site. No sequence mismatches were detected, confirming the absence of off-target editing at this site.
